# Proteomic Analysis of Urine from Youths Indulging in Gaming

**DOI:** 10.1101/2024.02.26.581984

**Authors:** Minhui Yang, Yuqing Liu, Ziyun Shen, Haitong Wang, Youhe Gao

**Affiliations:** Gene Engineering Drug and Biotechnology Beijing Key Laboratory, College of Life Sciences, Beijing Normal University, Beijing 100871, China

**Keywords:** urine, proteomics, addiction, video games

## Abstract

Video game addiction manifests as an escalating enthusiasm and uncontrolled use of digital games, yet there are no objective indicators for gaming addiction. This study employed mass spectrometry proteomics to analyze the proteomic differences in the urine of adolescents addicted to gaming compared to those who do not play video games. The study included 10 adolescents addicted to gaming and 9 non-gaming adolescents as a control group. The results showed that there were 125 significantly different proteins between the two groups. Among these, 11 proteins have been reported to change in the body after the intake of psychotropic drugs and are associated with addiction: Calmodulin, ATP synthase subunit alpha, ATP synthase subunit beta, Acid ceramidase, Tomoregulin-2, Calcitonin, Apolipoprotein E, Glyceraldehyde-3-phosphate dehydrogenase, Heat shock protein beta-1, CD63 antigen, Ephrin type-B receptor 4, Tomoregulin-2. Additionally, several proteins were found to interact with pathways related to addiction: Dickkopf-related protein 3, Nicastrin, Leucine-rich repeat neuronal protein 4, Cerebellin-4. Enriched biological pathways discovered include those related to nitric oxide synthase, amphetamine addiction, and numerous calcium ion pathways, all of which are associated with addiction. Moreover, through the analysis of differentially expressed proteins, we speculated about some proteins not yet fully studied, which might play a significant role in the mechanisms of addiction: Protein kinase C and casein kinase substrate in neurons protein, Cysteine-rich motor neuron 1 protein, Bone morphogenetic protein receptor type-2, Immunoglobulin superfamily member 8. In the analysis of urinary proteins in adolescents addicted to online gaming, we identified several proteins that have previously been reported in studies of drug addiction.

## 2. Introduction

Electronic game addiction, as an increasingly concerned issue in modern society, has attracted extensive research from experts and scholars worldwide. In recent years, with the rapid development of Internet technology and the booming rise of the electronic game industry, more and more people, especially teenagers, are addicted to electronic games, resulting in a series of psychological, social and physiological problems ^[1]^.

In the field of psychiatry, gaming addiction is seen as an impulse control disorder characterized by a strong desire for gaming, uncontrollable gaming behavior, loss of interest in other activities, and negative consequences of sustained gaming ^[2]^. These characteristics not only have a serious impact on personal health, but may also disrupt social relationships and quality of life. According to a global study, video game addiction is associated with various mental health issues, including depression, anxiety, and social dysfunction ^[3]^. In addition, neuroimaging studies have found that, similar to substance dependence, video game addiction can also lead to changes in brain structure and function ^[4,5]^. The formation of addiction to electronic games may be related to various factors, including personal psychological traits, social environment, and game design elements. Research has shown that reward systems, social interactions, and immersive experiences in games are key factors promoting addictive behavior ^[6]^.

Considering the severity and prevalence of electronic game addiction, understanding its addiction mechanisms and impacts is crucial for developing effective prevention and intervention strategies. Future research should continue to explore the neurobiological basis of game addiction and how to effectively address this challenge through psychological and social intervention measures.

Urine proteomics has shown significant potential in the discovery and analysis of biomarkers. Compared with other biological samples, urine has a unique advantage: it is not strictly regulated by physiological homeostasis mechanisms, and therefore can more sensitively reflect small biochemical changes in the body ^[7]^. In addition, the urine collection process is non-invasive and simple, making it an ideal source of biomarkers. Currently, numerous studies have confirmed that proteins in urine can serve as biomarkers for various neurological disorders in the brain, such as Parkinson’s syndrome ^[8]^, Alzheimer’s disease ^[9]^, depression ^[10]^, and autism ^[11]^. However, there is no research on game addiction in the field of urine proteomics. This article not only broadens our understanding of the mechanisms of electronic game addiction, but also hopes to help develop new therapies for this type of disease. Therefore, our technical roadmap for the study of urine proteomics in gaming addicted youth is shown in Figure 1.

**Figure 1:**
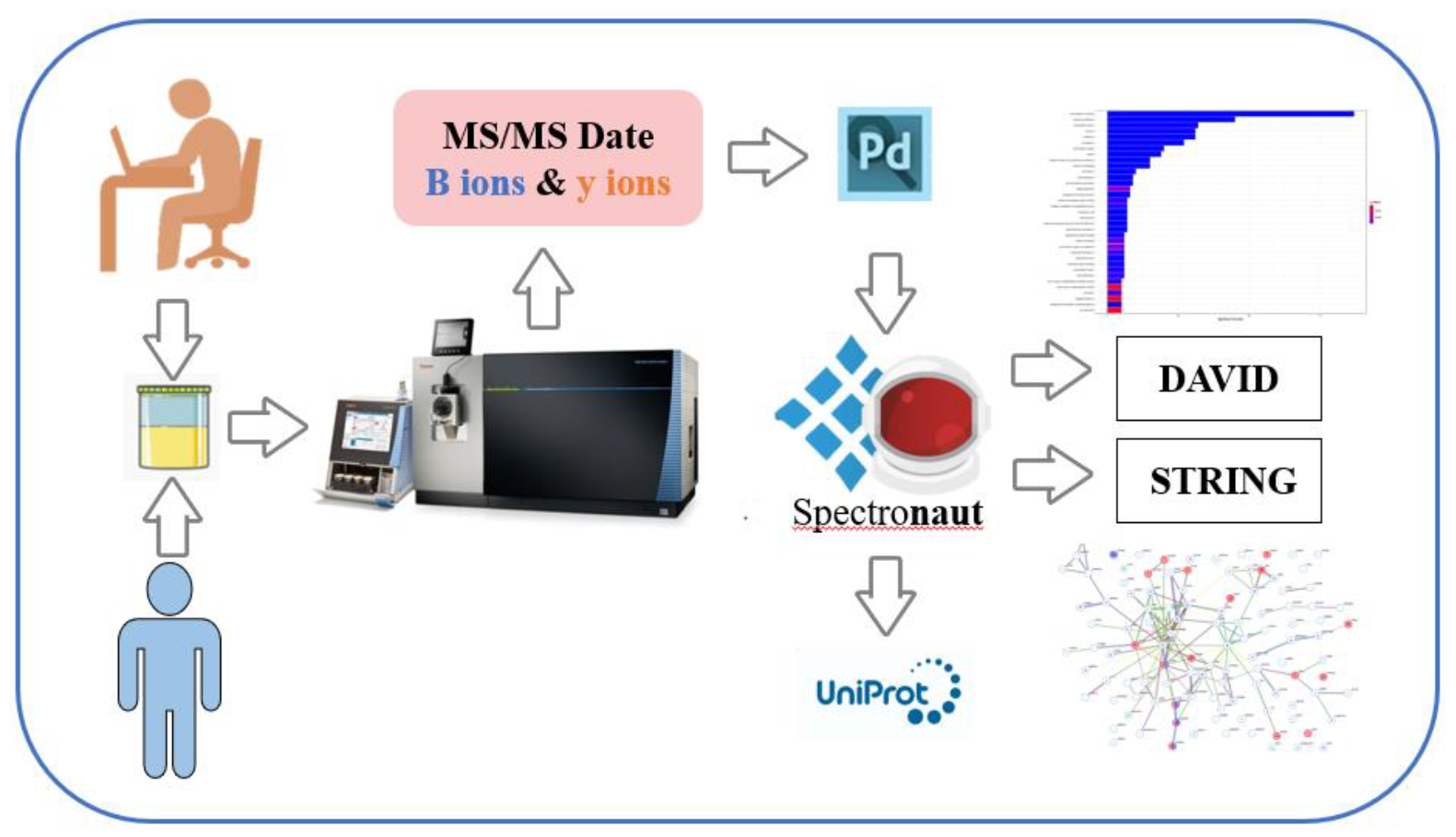
Technical route for urine proteomics analysis of gaming addicted youth

## 2. Materials and Methods

### 2.1 Sample Collection

This study collected urine from 10 gaming addicted youth (average age 22 years old) and 9 male non gaming youth (average age 24 years old), with sampling locations in internet cafes and participant homes. The sampling standard is in line with Young’s diagnostic criteria for moderate internet addiction, and all participants experience psychological discomfort when not playing games.

The daily gaming time is 6-9 hours. Volunteers have all signed informed consent books and provided them with detailed information about their research, including research objectives, methods, and processes. At the same time, personal information of volunteers is strictly kept confidential. There are no restrictions or requirements on the diet, medication, or other factors of the volunteers, and urine is collected and stored at -80 ° C. The sample information is detailed in Table 1.

**Table 1.** Subject Information.

| Subject number | Sex | age | Daily game duration |
| --- | --- | --- | --- |
| case 1 | 男 | 18 | 7-9h |
| case 2 | 男 | 20 | 8-9h |
| case 3 | 男 | 19 | 6-8h |
| case 4 | 男 | 22 | 8-9h |
| case 5 | 男 | 18 | 6-8h |
| case 6 | 男 | 22 | 8-9h |
| case 7 | 男 | 24 | 7-9h |
| case 8 | 男 | 19 | 6-8h |
| case 9 | 男 | 26 | 8-9h |
| case 10 | 男 | 20 | 6-8h |
| control 1 | 男 | 27 | 0h |
| control 2 | 男 | 24 | 0.5-1h |
| control 3 | 男 | 24 | 0.5-1h |
| control 4 | 男 | 20 | 0.5-1h |
| control 5 | 男 | 23 | 0.5-1h |
| control 6 | 男 | 27 | 0h |
| control 7 | 男 | 25 | 0.5-1h |
| control 8 | 男 | 28 | 0h |
| control 9 | 男 | 26 | 1h |

### 2.2 Treatment of Urine Samples

Firstly, centrifuge the urine sample at 12000 g for 30 minutes at 4 °C. Then, take 6 milliliters of urine and precipitate 15 milliliters of urine from each sample overnight with five times the volume of ethanol at -20 °C. After centrifugation at 12000 g, protein precipitates were obtained and dissolved in lysis buffer (8 mol/L urea, 2 mol/L thiourea, 50 mmol/L Tris, and 25 mmol/L dithiothreitol). Then, the Bradford analysis method was used for quantitative analysis of the supernatant.

Used trypsin digestion for a total of 100 μ G protein. Load the proteins in each sample into a 10 kDa filtration device. After washing twice with urea buffer and 25 mmol/L NH4HCO3 solution, reduce with 20 mmol/L dithiothreitol at 37 °C for 1 hour, alkylate with 50mmol/L iodoacetyl gel (IAA) in a dark environment for 45 minutes, then wash the sample with UA and NH4HCO3, and digest overnight with trypsin (enzyme to protein ratio 1:50) at 37 °C for 14-16 hours. Desalinate the digested peptides using the Oasis HLB assay kit, and then dry them using a freeze-drying machine. Dissolve the digested peptide substances in 0.1% formic acid, then dilute to 0.5 μg/μL.

### 2.3 LC-MS/MS tandem mass spectrometry analysis

The separated protein sample is subjected to ionization treatment and enters the mass spectrometer. Each sample is processed at a rate of 1 μ Analysis of peptide quality: Thermo EASY-nLC1200 chromatography system loaded onto the pre column and analysis column. Collect proteomic data using the Thermo Orbitrap Fusion Lumis mass spectrometry system.

### 2.4 Data processing and analysis

Each peptide sample was subjected to mass spectrometry data collection in DIA mode. The processing and analysis of mass spectrometry data are carried out using Spectraut X software. Search the database for the raw files (raw files) collected by DIA for each sample.

We apply the quantitative method of spectrogram number to screen for differential proteins. Screening criteria: Fold change ≥ 1.5 or ≤ 0.67, paired t-test analysis with P-value correction value<0.05. Subsequently, functional analysis of differential proteins was conducted through DAVID Functional Annotation Bioinformatics microarray Analysis and literature search in the Pubmed database.

## 3. Results and Discussion

### 3.1 Urine proteomic analysis

After processing the urine samples, LC-MS/MS tandem mass spectrometry analysis was performed on 19 protein samples. A total of 1205 proteins were identified (≥ 2 specific peptides, protein level FDR<1%).

A total of 125 differentially expressed proteins were identified by grouping 10 gaming addicted youth and 9 healthy samples, and 36 differentially expressed proteins were randomly generated, as shown in Table 1. The screening criteria for differential proteins are FC ≥ 1.5 or ≤ 0.67, with P<0.05. The specific information of differential proteins is listed in Table 2. By randomly matching the groups, an average of 36 differentially expressed proteins were obtained, which proves that the results of differentially expressed proteins are reliable and non randomly generated, as shown in Table 2.

**Table 2.** Random grouping.

| Fitter Criteria | Number of differential proteins | Grouping possibility | Average numbers of proteins with false random combinations | Reliability |
| --- | --- | --- | --- | --- |
| $FC > 1.5$ 或 $< 0.67$-$P < 0.05$ | 125 | 46189 | 36 | 71.2% |

A PubMed search was conducted on differentially expressed proteins that met the requirements (FC ≥ 1.5 or ≤ 0.67, P<0.05), and a total of 11 proteins were found to have been reported to be related to addiction, including ATP synthesis subunit alpha, ATP synthesis subunit beta, Glyceraldehyde-3 phase dehydrogenase, Acid ceramidase, Calcitonin, Apolipoprotein E, Calmodulin-1, Ephrin type-B receiver 4, Heat shock protein beta-1, CD63 antigen, Tomoregulin-2.

A series of differential proteins were discovered in the study, which exhibit unique changes under different drug addiction conditions. Taking ATP synthase subunit alpha as an example, as a key component of F-type ATP synthase ^[12]^, its expression level in the rat striatum decreased after long-term morphine treatment ^[13]^. The expression of ATP synthase subunit beta is also reduced in the nucleus accumbens of individuals with cocaine overdose ^[14,15]^. In the observation of morphine exposure for 10 days and cessation of medication for 20 days, another key protein - Glyceraldehyde-3-phosphate dehydrogenase also showed significant expression changes ^[16]^. Wenchao Li and colleagues used high-throughput chain specific RNA sequencing to investigate how cocaine affects gene expression in the insula of mice. Their GO enrichment analysis revealed the regulatory effect of cocaine on synaptic transmission of ceramidase activity ^[17]^, and in this experimental group, the differential protein Acid ceramidase was 2.5 times higher than that in the control group. In addition, studies have shown that serum levels of calcitonin significantly increase after cocaine withdrawal^[18]^.

Research on methamphetamine addiction has revealed that acute exposure to the drug increases the expression level of Apolipoprotein E (ApoE) protein in the brain parenchyma. This elevation activates ApoE receptor-2 on the cerebral capillaries and affects the expression of LTP ^[19]^. MICHELHAUGH S K’s research further indicates that methamphetamine can significantly increase the content of calmodulin and messenger RNA levels in rat brain regions ^[18,19]^. A comparative study between methamphetamine abstainers (WMA) and healthy controls (HC) found significant differences in the expression of Ephrin type-B receptor 2 (EPHB2) ^[20]^. This discovery suggests that EPHB2 may play a role in regulating the glutamate metabolism pathway. In the study of the effects of methamphetamine withdrawal on the activation of the brain stress system and cardiac sympathetic nervous pathways, it was found that methamphetamine withdrawal increased the level of Heat shock protein beta-1 in the heart ^[21]^. In addition, compared with the healthy control group, the expression of CD63 antigen in platelets of heroin, cocaine, and cannabis addicts was significantly reduced ^[22]^.

Tomoregulin-2 can promote ERK1/2 phosphorylation, and extracellular signal regulated kinase (ERK) has been shown to be activated by opioid drugs and functionally associated with addiction ^[23]^. The formation of a complex between Tomoregulin-1 and addicosin (a protein highly upregulated in the amygdala of morphine treated mice) in the same family affects the distribution of Tomoregulin-1 within cells and regulates cell migration, demonstrating that Tomoregulin-1 is a novel addicosin related factor ^[24]^. Tomoregulin-1 and Tomoregulin-2 are both proteins belonging to the TMEFF family, and Tomoregulin-2 has been shown to be highly expressed in all brain regions except the pituitary gland, expressed in the amygdala and corpus callosum. It is a survival factor for hippocampal neurons and midbrain neurons ^[23]^, and exists in Alzheimer’s disease plaques ^[25]^.

It is worth mentioning that Tomoregulin-2 may inhibit Bone Morphogenic Proteins (BMP) signaling during neural pattern formation ^[26]^. BMP plays a crucial role in the central nervous system, especially in terms of neural plasticity. They play a role in multiple stages of central nervous system development, including the formation and patterning of the brain and spinal cord. In the adult brain, BMP is detected in regions related to neural plasticity and has been shown to regulate neurogenesis, glial cell formation, and synaptic and dendritic plasticity ^[27, 28]^. These findings indicate that BMP is crucial for the dynamic adjustment of brain structure and function.

It is interesting that we also found ATP synthase subunits in differential proteins α And ATP synthase subunits β. ATP synthase is a key intracellular enzyme whose main function is to catalyze the synthesis of adenosine triphosphate (ATP). This complex enzyme structure is composed of multiple subunits, among which subunits α (alpha) and subunits β (beta) plays a core role in its F1 section and is crucial for the ATP synthesis process ^[29]^.

Yaji α, As the main structural component of ATP synthase, located in the F1 region, each ATP synthase complex typically contains three α Yaji. although α The subunit does not directly participate in ATP synthesis, but it plays a crucial role in maintaining the structural stability and functional normalization of the entire enzyme complex. It is related to β The alternating arrangement of subunits forms a circular structure, providing necessary structural support for the catalytic center of ATP synthesis ^[30]^. Yaji β, Another key component located in the F1 section, which also includes three β Yaji. Related to α Different subunits, β The subunit directly participates in the synthesis of ATP and contains active catalytic sites. During the operation of ATP synthase, β The subunit undergoes various conformational changes, which are crucial for the synthesis and release of ATP. In addition to ATP synthase, Apolipoprotein E is closely related to energy metabolism, and the changes in other proteins are also worth exploring.

Immunoglobulin superfamily member 8 (IgSF8) interacts with integrin, an extracellular matrix receptor that mediates biochemical and mechanical bidirectional signals between the extracellular and intracellular environments due to conformational changes. In the brain, they exist in neurons and glial cells, playing important roles in several aspects of brain development and function, such as cell migration, axonal guidance, synaptogenesis, synaptic plasticity, and neuroinflammation. There are also reports that integrins are related to addiction ^[32]^. In addition, Immunoglobulin superfamily member 21 interacts with neurotransmitters ^[33]^, which are also associated with addictive behavior ^[34]^. Based on these findings, we can speculate that IgSF8 may play an important role in the process of addiction.

Dickkopf related protein 3 locally inhibits the process of Wnt regulation ^[35]^, and the Wnt pathway plays an important role in the development of the midbrain dopaminergic system, where dopaminergic neurons affect addiction and other neurological disorders ^[36]^. In addition, Uniprot analyzed the role of Nicastin in the regulation of the Wnt signaling pathway and downstream processes through similarity analysis.

Protein kinase C and casein kinase substrate in neurons protein 3 were only 0.13 times higher than the control group, and they play a role in endocytosis and regulate the internalization of plasma membrane proteins ^[37]^. In adulthood, PACSIN1 has been extensively studied in the brain and has been shown to regulate neuromorphogenesis, receptor transport, and synaptic plasticity ^[38]^.

Our research suggests that there may be similar biological mechanisms between game addiction and drug addiction. This is supported by the common changes in addictive behavior related proteins and neurotransmitter pathways. This similarity suggests that although the objects of addiction are different (such as games or drugs), they may follow similar patterns in biology.

### 3.2 Enrichment pathway analysis

Enrichment of pathways related to amphetamine addiction, response to corticosterone, calcium ion, and nitric oxide synthase was analyzed through DAVID Functional Annotation Bioinformatics microarray Analysis, as shown in Tables 3 and 4.

**Table 3.**
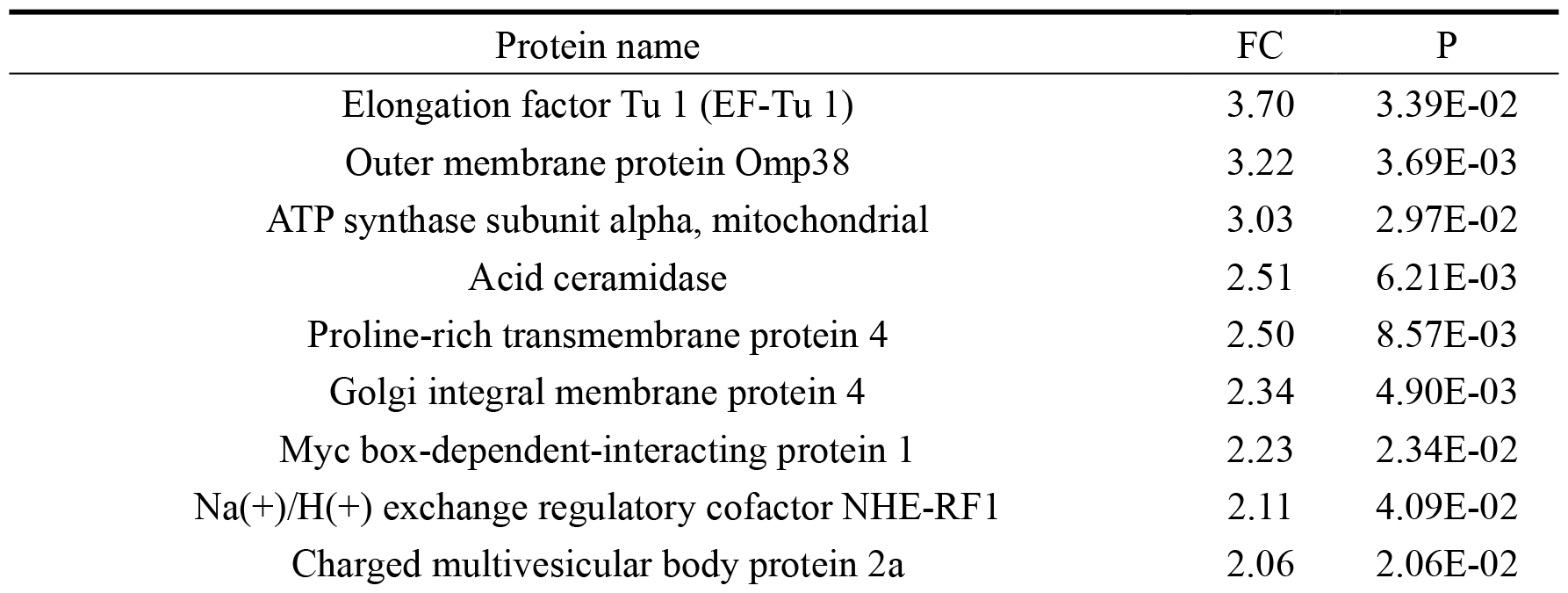
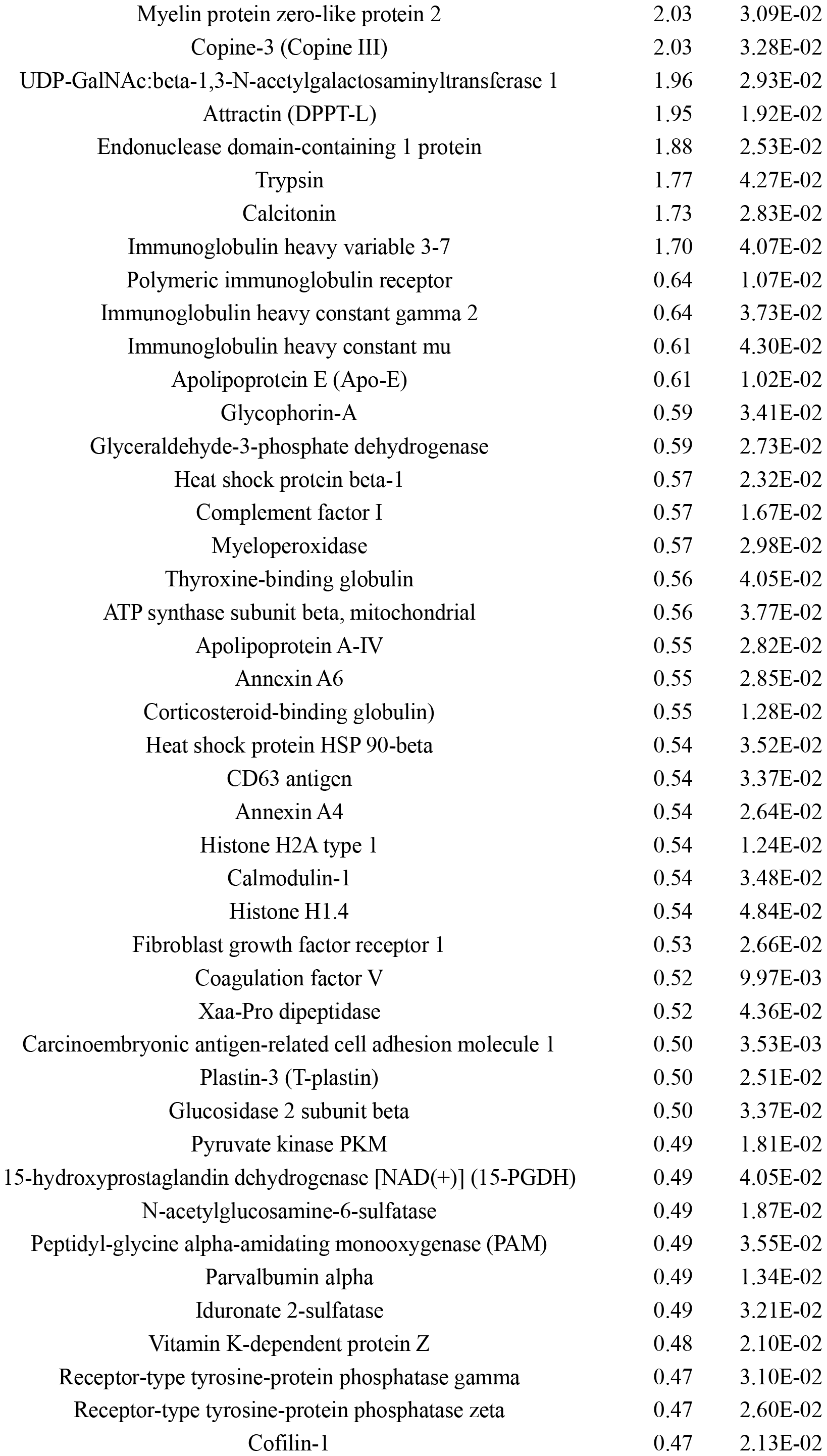
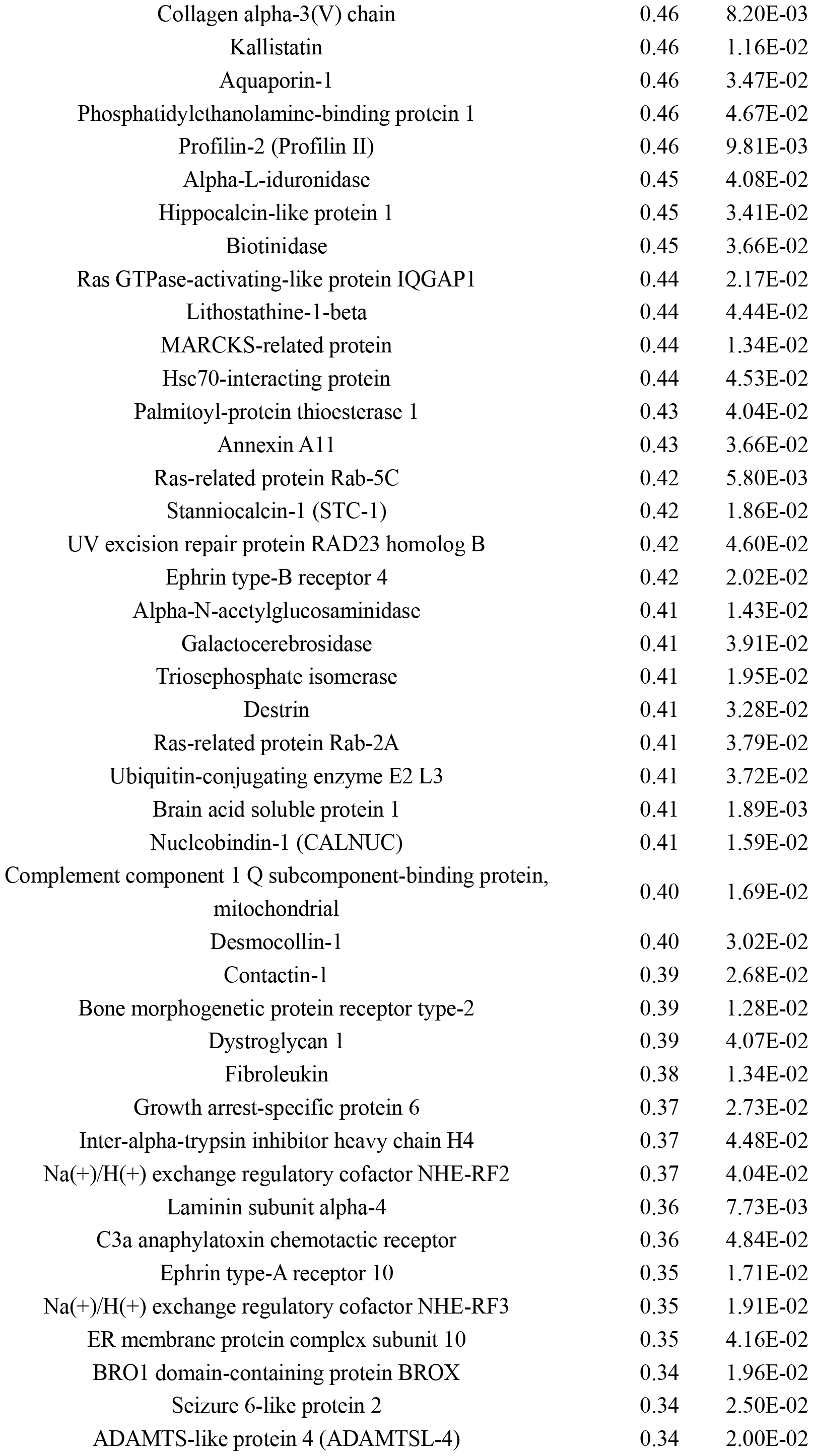
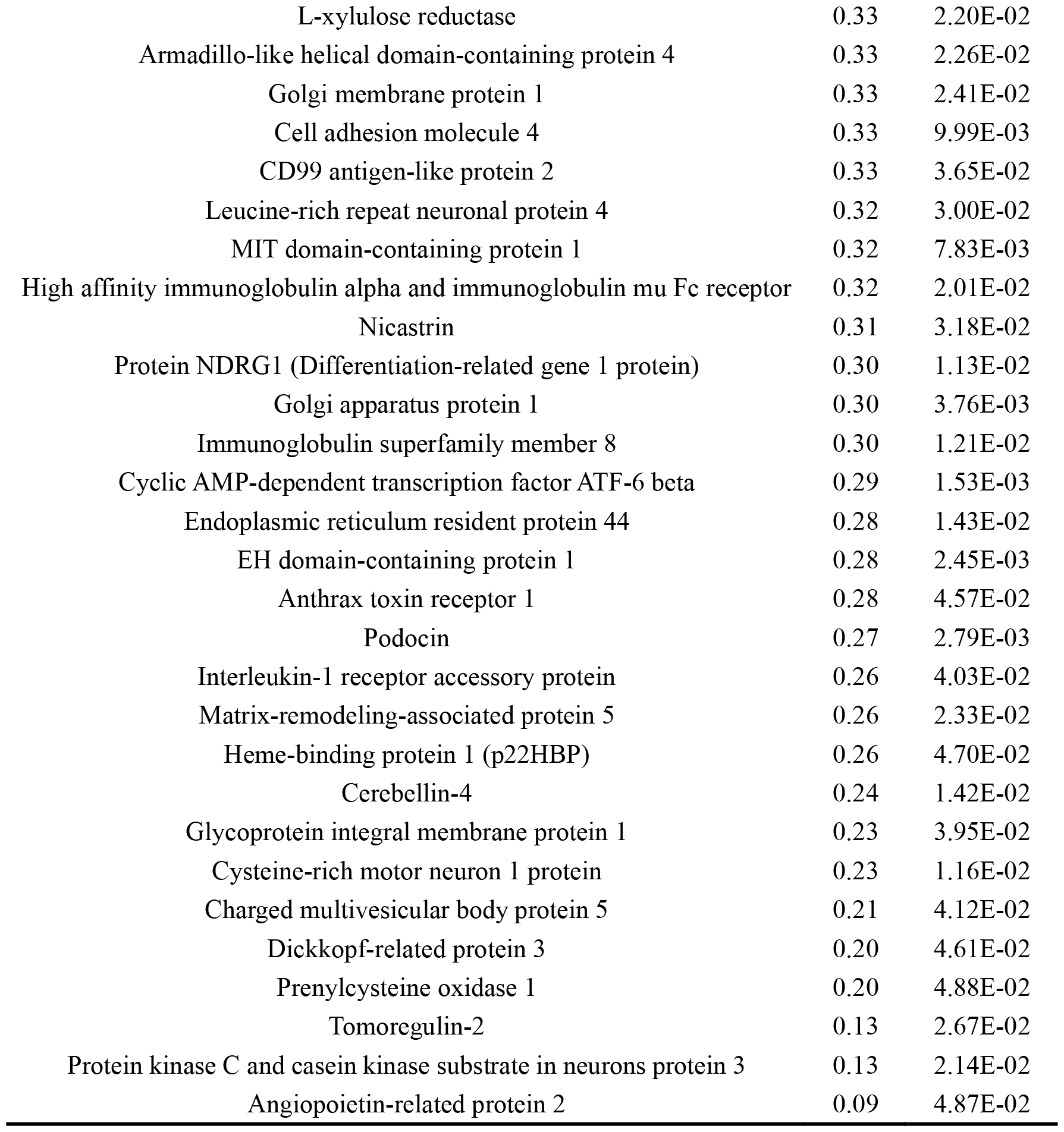
Differential Proteins.

**Table 4.**
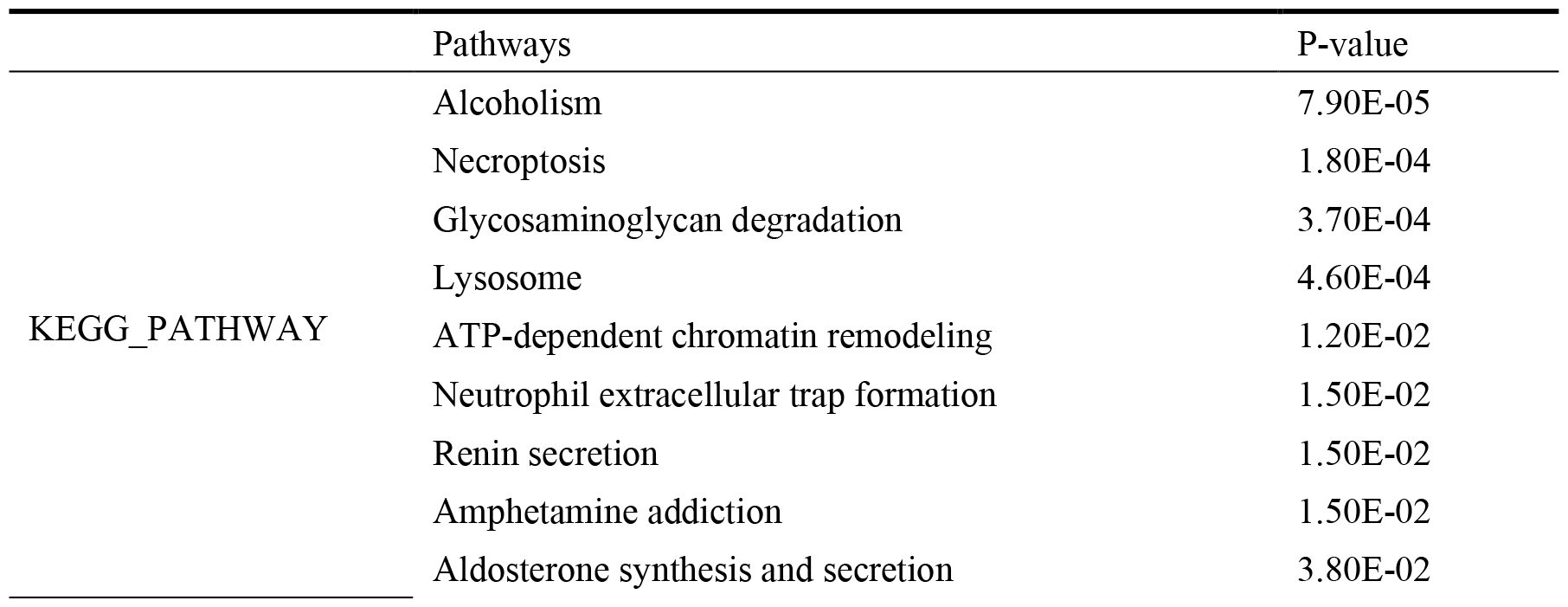
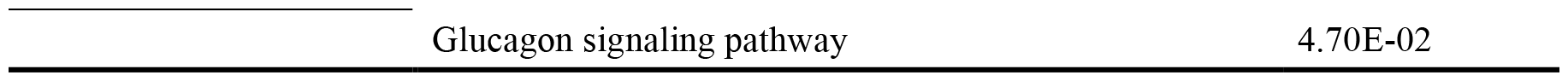
KEGG Enrichment Pathways.

**Table 5.**
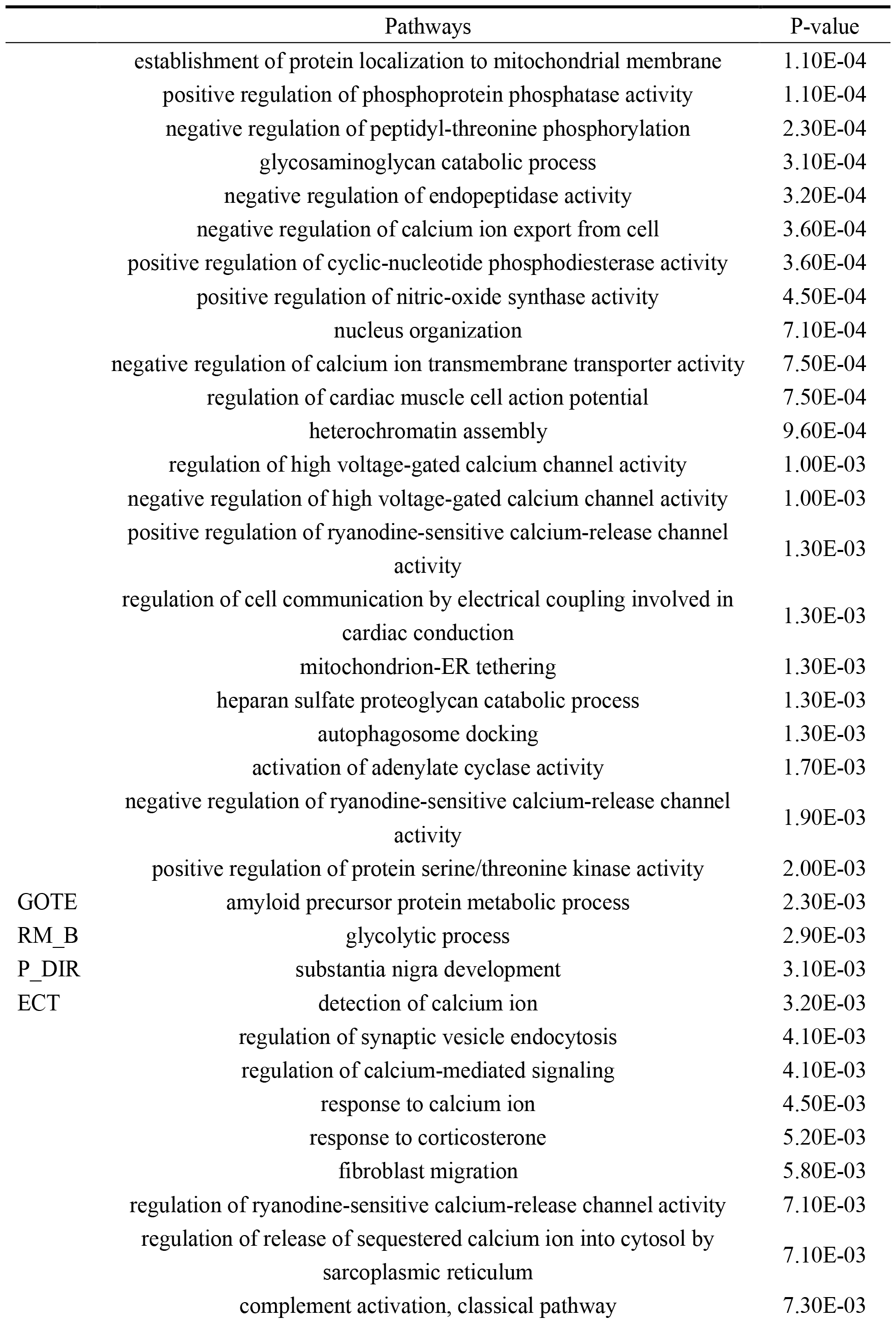
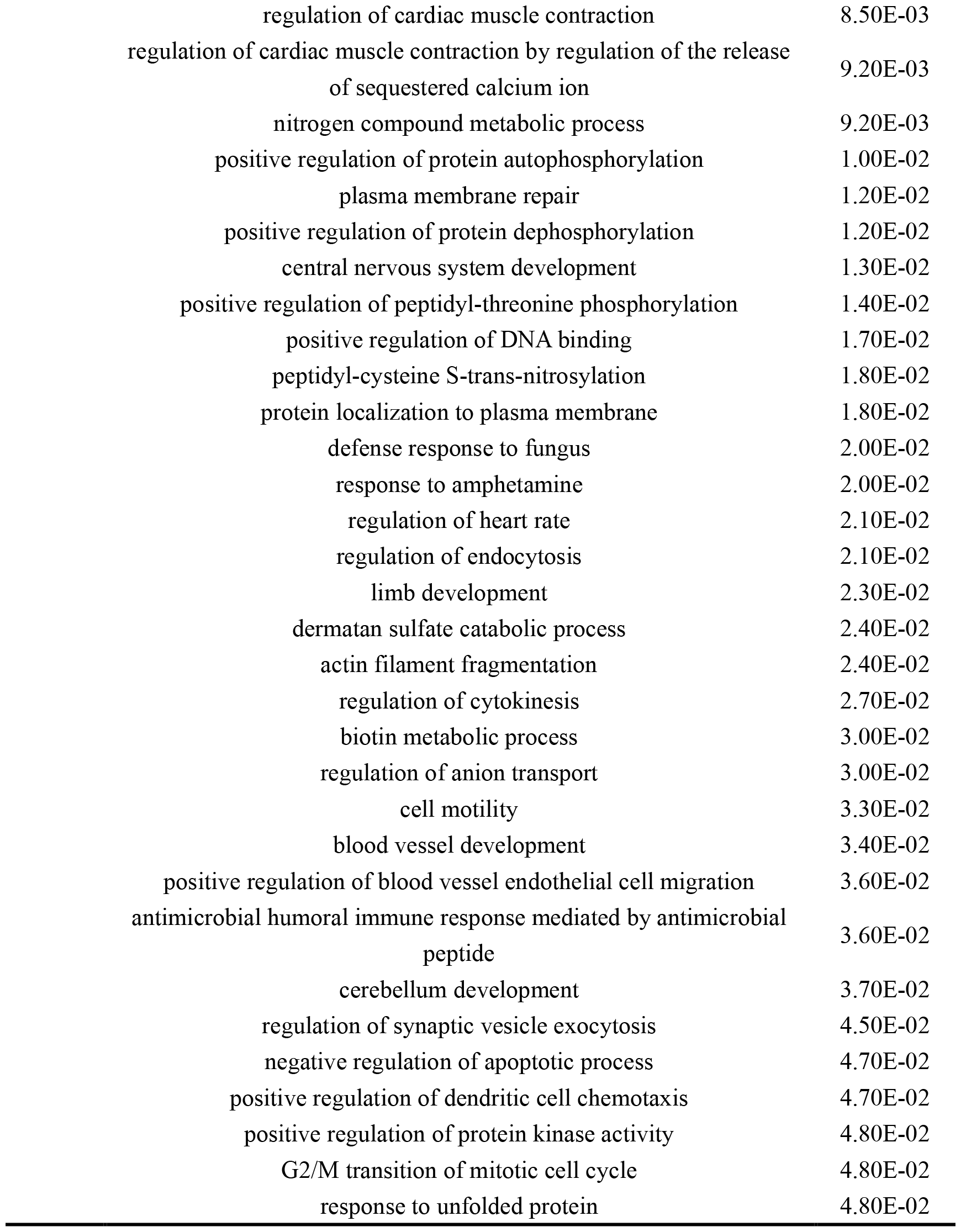
GO Enrichment Pathways.

|  | Pathways | P-value |
| --- | --- | --- |
|  | establishment of protein localization to mitochondrial membrane | 1.10E-04 |
|  | positive regulation of phosphoprotein phosphatase activity | 1.10E-04 |
|  | negative regulation of peptidyl-threonine phosphorylation | 2.30E-04 |
|  | glycosaminoglycan catabolic process | 3.10E-04 |
|  | negative regulation of endopeptidase activity | 3.20E-04 |
|  | negative regulation of calcium ion export from cell | 3.60E-04 |
|  | positive regulation of cyclic-nucleotide phosphodiesterase activity | 3.60E-04 |
|  | positive regulation of nitric-oxide synthase activity | 4.50E-04 |
|  | nucleus organization | 7.10E-04 |
|  | negative regulation of calcium ion transmembrane transporter activity | 7.50E-04 |
|  | regulation of cardiac muscle cell action potential | 7.50E-04 |
|  | heterochromatin assembly | 9.60E-04 |
|  | regulation of high voltage-gated calcium channel activity | 1.00E-03 |
|  | negative regulation of high voltage-gated calcium channel activity | 1.00E-03 |
|  | positive regulation of ryanodine-sensitive calcium-release channel activity | 1.30E-03 |
|  | regulation of cell communication by electrical coupling involved in cardiac conduction | 1.30E-03 |
|  | mitochondrion-ER tethering | 1.30E-03 |
|  | heparan sulfate proteoglycan catabolic process | 1.30E-03 |
|  | autophagosome docking | 1.30E-03 |
|  | activation of adenylate cyclase activity | 1.70E-03 |
|  | negative regulation of ryanodine-sensitive calcium-release channel activity | 1.90E-03 |
|  | positive regulation of protein serine/threonine kinase activity | 2.00E-03 |
| GOTE | amyloid precursor protein metabolic process | 2.30E-03 |
| RM_B | glycolytic process | 2.90E-03 |
| P_DIR | substantia nigra development | 3.10E-03 |
| ECT | detection of calcium ion | 3.20E-03 |
|  | regulation of synaptic vesicle endocytosis | 4.10E-03 |
|  | regulation of calcium-mediated signaling | 4.10E-03 |
|  | response to calcium ion | 4.50E-03 |
|  | response to corticosterone | 5.20E-03 |
|  | fibroblast migration | 5.80E-03 |
|  | regulation of ryanodine-sensitive calcium-release channel activity | 7.10E-03 |
|  | regulation of release of sequestered calcium ion into cytosol by sarcoplasmic reticulum | 7.10E-03 |
|  | complement activation, classical pathway | 7.30E-03 |

|  |  |
| --- | --- |
| regulation of cardiac muscle contraction | 8.50E-03 |
| regulation of cardiac muscle contraction by regulation of the release<br>of sequestered calcium ion | 9.20E-03 |
| nitrogen compound metabolic process | 9.20E-03 |
| positive regulation of protein autophosphorylation | 1.00E-02 |
| plasma membrane repair | 1.20E-02 |
| positive regulation of protein dephosphorylation | 1.20E-02 |
| central nervous system development | 1.30E-02 |
| positive regulation of peptidyl-threonine phosphorylation | 1.40E-02 |
| positive regulation of DNA binding | 1.70E-02 |
| peptidyl-cysteine S-trans-nitrosylation | 1.80E-02 |
| protein localization to plasma membrane | 1.80E-02 |
| defense response to fungus | 2.00E-02 |
| response to amphetamine | 2.00E-02 |
| regulation of heart rate | 2.10E-02 |
| regulation of endocytosis | 2.10E-02 |
| limb development | 2.30E-02 |
| dermatan sulfate catabolic process | 2.40E-02 |
| actin filament fragmentation | 2.40E-02 |
| regulation of cytokinesis | 2.70E-02 |
| biotin metabolic process | 3.00E-02 |
| regulation of anion transport | 3.00E-02 |
| cell motility | 3.30E-02 |
| blood vessel development | 3.40E-02 |
| positive regulation of blood vessel endothelial cell migration | 3.60E-02 |
| antimicrobial humoral immune response mediated by antimicrobial<br>peptide | 3.60E-02 |
| cerebellum development | 3.70E-02 |
| regulation of synaptic vesicle exocytosis | 4.50E-02 |
| negative regulation of apoptotic process | 4.70E-02 |
| positive regulation of dendritic cell chemotaxis | 4.70E-02 |
| positive regulation of protein kinase activity | 4.80E-02 |
| G2/M transition of mitotic cell cycle | 4.80E-02 |
| response to unfolded protein | 4.80E-02 |

By using the Gene Ontology (GO) analysis method, we discovered a remarkable phenomenon: we found many calcium ion related pathways in the pathways, including negative regulation of cellular calcium ion output, negative regulation of calcium ion transmembrane transport protein activity, regulation of high-pressure gated calcium channel activity, positive regulation of Ryanodine sensitive calcium release channel activity, calcium mediated signaling regulation, calcium ion response, and regulation of fixed calcium ion release to the cytoplasm by the sarcoplasmic reticulum.

Long term potential (LTP) is a crucial neurobiological process that leads to enhanced synaptic connections and plays a crucial role in learning and memory formation. A key triggering factor of LTP is the increase in calcium ion (Ca2+) concentration in the postsynaptic region. This increase in Ca2+concentration activates a series of complex intracellular signaling pathways. Specifically, the increase of Ca2+triggers the generation of cyclic adenosine monophosphate (cAMP), which is achieved by activating adenosine monophosphate cyclase. Subsequently, cAMP activates protein kinase A (PKA), which in turn leads to a sustained increase in the amount of glutamate released by nerve endings at each action potential. LTP also plays an important role in addictive neural adaptation. Research has shown that drug abuse can induce LTP in the reward circuit of the brain, especially in the ventral tegmental area (VTA), a key area closely related to the development of addiction. It is worth noting that presynaptic LTP is mainly triggered by a dependent increase in Ca2+activity in presynaptic terminals ^[39, 40]^.

In the differential proteins we studied, we particularly paid attention to Cerebellin-4 (Cbln4). The high expression of Cbln4 in the olfactory cortex is particularly important, as it plays a crucial role in the long-term synaptic enhancement (LTP) from the olfactory cortex to the dentate gyrus. Specifically, the binding of presynaptic Cbln4 to postsynaptic neogenin-1 is necessary for the LTP from the olfactory cortex to the dentate gyrus synapse ^[41]^. Based on this discovery, we speculate that Cerebellin-4 may also play a key role in the addiction process. In addition, Leucine rich repeat neural protein 4 plays a role in hippocampal dependent persistent memory ^[42]^, which may play an important role in addiction.

We have also enriched the positive regulation of nitric oxide synthase activity in the pathway, and nitric oxide, as an important neurotransmitter, plays a crucial role in the central nervous system. Numerous studies have suggested that nitric oxide may play an important role in drug addiction, including dependence on opioids, alcohol, stimulants, and nicotine. Of particular note, inhibitors targeting nitric oxide synthase have been found to effectively regulate withdrawal symptoms caused by these addictive substances ^[43]^.

It is worth noting that there is a significant correlation between game addiction and the amphetamine addiction pathway in the protein expression pattern of brain tissue. This discovery reveals that gaming addiction and drug addiction may affect the same neurobiological pathways, which have been extensively studied in more widely recognized addictive substances such as amphetamines. Our results provide a new perspective on how electronic games affect the brain at the molecular level and valuable clues for further research on how caffeine and other stimulants affect brain function and addictive behavior.

These findings reveal the sensitivity of urine in reflecting addiction status, providing a new perspective for understanding the essence of addiction. The article mentions changes in calcium ions, nitric oxide synthase, ATP synthase, etc., which have been proven by many studies to be related to addiction. It is worth noting that other proteins and pathways that have not been studied in the field of addiction may play important roles in addiction mechanisms, and this article provides reliable clues.

## 4. Conclusion

Teenagers addicted to online games exhibit significant differences in urine protein composition compared to non gamers. Among the differentially proteins, we identified multiple proteins previously reported in drug addiction studies.

## References

[1] Boz C, Dinc M. Examination of game addiction studies conducted in Turkey: A systematic review study [J]. Front Psychiatry, 2023, 14: 1014621.

[2] Yalcin Irmak A, Erdogan S. [Digital Game Addiction Among Adolescents and Younger Adults: A Current Overview] [J]. Turk Psikiyatri Derg, 2016, 27(2): 0.

[3] Gentile D A, Choo H, Liau A, et al. Pathological video game use among youths: a two-year longitudinal study [J]. Pediatrics, 2011, 127(2): e319–29.

[4] Kuhn S, Gallinat J. Brain structure and functional connectivity associated with pornography consumption: the brain on porn [J]. JAMA Psychiatry, 2014, 71(7): 827–34.

[5] Mohammad S, Jan R A, Alsaedi S L. Symptoms, Mechanisms, and Treatments of Video Game Addiction [J]. Cureus, 2023, 15(3): e36957.

[6] Kuss D J, Griffiths M D. Internet and gaming addiction: a systematic literature review of neuroimaging studies [J]. Brain Sci, 2012, 2(3): 347–74.

[7] Wu J, Li X, Zhao M, et al. Early Detection of Urinary Proteome Biomarkers for Effective Early Treatment of Pulmonary Fibrosis in a Rat Model [J]. Proteomics Clin Appl, 2017, 11(11-12).

[8] Virreira Winter S, Karayel O, Strauss M T, et al. Urinary proteome profiling for stratifying patients with familial Parkinson’s disease [J]. EMBO Mol Med, 2021, 13(3): e13257.

[9] Watanabe Y, Hirao Y, Kasuga K, et al. Urinary Apolipoprotein C3 Is a Potential Biomarker for Alzheimer’s Disease [J]. Dement Geriatr Cogn Dis Extra, 2020, 10(3): 94–104.

[10] Huan Y, Wei J, Zhou J, et al. Label-Free Liquid Chromatography-Mass Spectrometry Proteomic Analysis of the Urinary Proteome for Measuring the Escitalopram Treatment Response From Major Depressive Disorder [J]. Front Psychiatry, 2021, 12: 700149.

[11] Wang Y, Zhang J, Song W, et al. A proteomic analysis of urine biomarkers in autism spectrum disorder [J]. J Proteomics, 2021, 242: 104259.

[12] Lee J H, Garboczi D N, Thomas P J, et al. Mitochondrial ATP synthase. cDNA cloning, amino acid sequence, overexpression, and properties of the rat liver alpha subunit [J]. J Biol Chem, 1990, 265(8): 4664–9.

[13] Bierczynska-Krzysik A, Pradeep John J P, Silberring J, et al. Proteomic analysis of rat cerebral cortex, hippocampus and striatum after exposure to morphine [J]. Int J Mol Med, 2006, 18(4): 775–84.

[14] Luo Y, Liao C, Chen L, et al. Heroin Addiction Induces Axonal Transport Dysfunction in the Brain Detected by In Vivo MRI [J]. Neurotox Res, 2022, 40(4): 1070–85.

[15] Tannu N, Mash D C, Hemby S E. Cytosolic proteomic alterations in the nucleus accumbens of cocaine overdose victims [J]. Mol Psychiatry, 2007, 12(1): 55–73.

[16] Ujcikova H, Cechova K, Jagr M, et al. Proteomic analysis of protein composition of rat hippocampus exposed to morphine for 10 days; comparison with animals after 20 days of morphine withdrawal [J]. PLoS One, 2020, 15(4): e0231721.

[17] Li W, Zhang C, Wang Y Y, et al. Alterations of RNAs in the insula related to cocaine-induced condition place preference in adolescent mice [J]. Biochem Biophys Res Commun, 2022, 621: 109–15.

[18] Spagnolli W, Torboli P, Mattarei M, et al. Calcitonin and prolactin serum levels in heroin addicts: study on a methadone treated group [J]. Drug Alcohol Depend, 1987, 20(2): 143–8.

[19] Michelhaugh S K, Gnegy M E. Differential regulation of calmodulin content and calmodulin messenger RNA levels by acute and repeated, intermittent amphetamine in dopaminergic terminal and midbrain areas [J]. Neuroscience, 2000, 98(2): 275–85.

[20] Peng S, Su H, Chen T, et al. The Potential Regulatory Network of Glutamate Metabolic Pathway Disturbance in Chinese Han Withdrawal Methamphetamine Abusers [J]. Front Genet, 2021, 12: 653443.

[21] Garcia-Carmona J A, Georgiou P, Zanos P, et al. Methamphetamine withdrawal induces activation of CRF neurons in the brain stress system in parallel with an increased activity of cardiac sympathetic pathways [J]. Naunyn Schmiedebergs Arch Pharmacol, 2018, 391(4): 423–34.

[22] Bell C R, Horowitz E D, Oshrine B R, et al. Profound inhibition of GPIb, GPIIb/IIIa, PECAM-1, CD63, and CD107 in a chronic drug addict: selecting controls for platelet flow cytometry in the inner city hospital [J]. Thromb Res, 2001, 101(3): 217–8.

[23] Horie M, Mitsumoto Y, Kyushiki H, et al. Identification and characterization of TMEFF2, a novel survival factor for hippocampal and mesencephalic neurons [J]. Genomics, 2000, 67(2): 146–52.

[24] Arano T, Fujisaki S, Ikemoto M J. Identification of tomoregulin-1 as a novel addicsin-associated factor [J]. Neurochem Int, 2014, 71: 22–35.

[25] Siegel D A, Davies P, Dobrenis K, et al. Tomoregulin-2 is found extensively in plaques in Alzheimer’s disease brain [J]. J Neurochem, 2006, 98(1): 34–44.

[26] Liu A, Niswander L A. Bone morphogenetic protein signalling and vertebrate nervous system development [J]. Nat Rev Neurosci, 2005, 6(12): 945–54.

[27] Jensen G S, Leon-Palmer N E, Townsend K L. Bone morphogenetic proteins (BMPs) in the central regulation of energy balance and adult neural plasticity [J]. Metabolism, 2021, 123: 154837.

[28] Manzari-Tavakoli A, Babajani A, Farjoo M H, et al. The Cross-Talks Among Bone Morphogenetic Protein (BMP) Signaling and Other Prominent Pathways Involved in Neural Differentiation [J]. Front Mol Neurosci, 2022, 15: 827275.

[29] Walker J E. The ATP synthase: the understood, the uncertain and the unknown [J]. Biochem Soc Trans, 2013, 41(1): 1–16.

[30] Junge W, Nelson N. ATP synthase [J]. Annu Rev Biochem, 2015, 84: 631–57.

[31] Stipp C S, Kolesnikova T V, Hemler M E. EWI-2 regulates alpha3beta1 integrin-dependent cell functions on laminin-5 [J]. J Cell Biol, 2003, 163(5): 1167–77.

[32] Jaudon F, Thalhammer A, Cingolani L A. Integrin adhesion in brain assembly: From molecular structure to neuropsychiatric disorders [J]. Eur J Neurosci, 2021, 53(12): 3831–50.

[33] Tanabe Y, Naito Y, Vasuta C, et al. IgSF21 promotes differentiation of inhibitory synapses via binding to neurexin2alpha [J]. Nat Commun, 2017, 8(1): 408.

[34] Kasem E, Kurihara T, Tabuchi K. Neurexins and neuropsychiatric disorders [J]. Neurosci Res, 2018, 127: 53–60.

[35] Yu C, Seaton M, Letendre S, et al. Plasma dickkopf-related protein 1, an antagonist of the Wnt pathway, is associated with HIV-associated neurocognitive impairment [J]. AIDS, 2017, 31(10): 1379–85.

[36] Alves Dos Santos M T, Smidt M P. En1 and Wnt signaling in midbrain dopaminergic neuronal development [J]. Neural Dev, 2011, 6: 23.

[37] Modregger J, Ritter B, Witter B, et al. All three PACSIN isoforms bind to endocytic proteins and inhibit endocytosis [J]. J Cell Sci, 2000, 113 Pt 24: 4511–21.

[38] Dumont V, Lehtonen S. PACSIN proteins in vivo: Roles in development and physiology [J]. Acta Physiol (Oxf), 2022, 234(3): e13783.

[39] Malenka R C, Bear M F. LTP and LTD: an embarrassment of riches [J]. Neuron, 2004, 44(1): 5–21.

[40] Nicoll R A, Schmitz D. Synaptic plasticity at hippocampal mossy fibre synapses [J]. Nat Rev Neurosci, 2005, 6(11): 863–76.

[41] Liakath-Ali K, Polepalli J S, Lee S J, et al. Transsynaptic cerebellin 4-neogenin 1 signaling mediates LTP in the mouse dentate gyrus [J]. Proc Natl Acad Sci U S A, 2022, 119(20): e2123421119.

[42] Bando T, Sekine K, Kobayashi S, et al. Neuronal leucine-rich repeat protein 4 functions in hippocampus-dependent long-lasting memory [J]. Mol Cell Biol, 2005, 25(10): 4166–75.

[43] Tayfun Uzbay I, Oglesby M W. Nitric oxide and substance dependence [J]. Neurosci Biobehav Rev, 2001, 25(1): 43–52.

